# Determination of *Pseudomonas aeruginosa* MexXY-OprM substrate profile in a major efflux knockout system reveals distinct antibiotic substrate classes

**DOI:** 10.1101/2024.10.18.619101

**Authors:** Logan G. Kavanaugh, Shraddha M. Hariharan, Graeme L. Conn

**Affiliations:** Department of Biochemistry, Emory University School of Medicine, Atlanta, GA; Microbiology and Molecular Genetics Graduate Program, Emory University, Atlanta, GA; Emory Antibiotic Resistance Center, Emory University, Atlanta, GA

**Keywords:** Efflux, *Pseudomonas*, antimicrobial susceptibility, antimicrobial resistance, resistance-nodulation-division (RND)

## Abstract

Defining the substrates of Resistance-Nodulation-Division (RND) systems is often complicated by the presence of multiple systems in a single bacterial species. Using a major efflux knockout strain, we developed a *Pseudomonas aeruginosa* system that enables controlled expression of MexXY-OprM in the absence of background efflux complexes with overlapping substrate profiles. This system identified three groups of potential substrates: substrates, partial substrates, and nonsubstrates, and defined trimethoprim as a new substrate for MexXY-OprM.

---

*Pseudomonas aeruginosa* is a Gram-negative, multidrug resistant (MDR) opportunistic pathogen associated with acute and chronic infections, linked to ventilators and catheters in nosocomial settings and, for example, localized in the lungs of individuals with cystic fibrosis and chronic obstructive pulmonary disease, respectively^1-3^. In 2019, MDR *P. aeruginosa* contributed to >300,000 deaths worldwide^1^, with the occurrence of hospital-associated infections increasing by 32% in 2020 due to the COVID-19 pandemic driving longer hospital stays and increasing the prevalence of secondary infections^2^. A major contributor to resistance in *P. aeruginosa* is active removal of antimicrobials via efflux, particularly by members of the <u>R</u>esistance-<u>N</u>odulation-<u>D</u>ivision (RND) efflux family^4-7^.

RND efflux pumps are tripartite complexes comprising a trimeric outer membrane protein (OMP), a hexameric periplasmic adaptor protein (PAP), and a trimeric inner membrane (IM) transporter^8^, e.g., OprM, MexX and MexY, respectively (**Fig. 1A**). MexXY-OprM is recognized for its unique ability in *P. aeruginosa* to efflux aminoglycosides, but its substrate profile is also proposed to include ethidium bromide (EtBr), tetracyclines, macrolides, chloramphenicol (Chl), and some β-lactams^5, 9-16^. Prior studies have examined substrate preference in a limited efflux knockout background (e.g., Δ*mexAB*, Δ*mexC*D, and/or Δ*mexXY*), through heterologous expression (e.g., in *E. coli*), and/or with uncontrolled induction systems (e.g., knockout of the native *mexXY* repressor, MexZ)^5, 9-16^. As such, these systems do not account for compensating effects from the remaining RND pumps and determining the specific contribution of MexXY can be challenging.

**Fig. 1.**
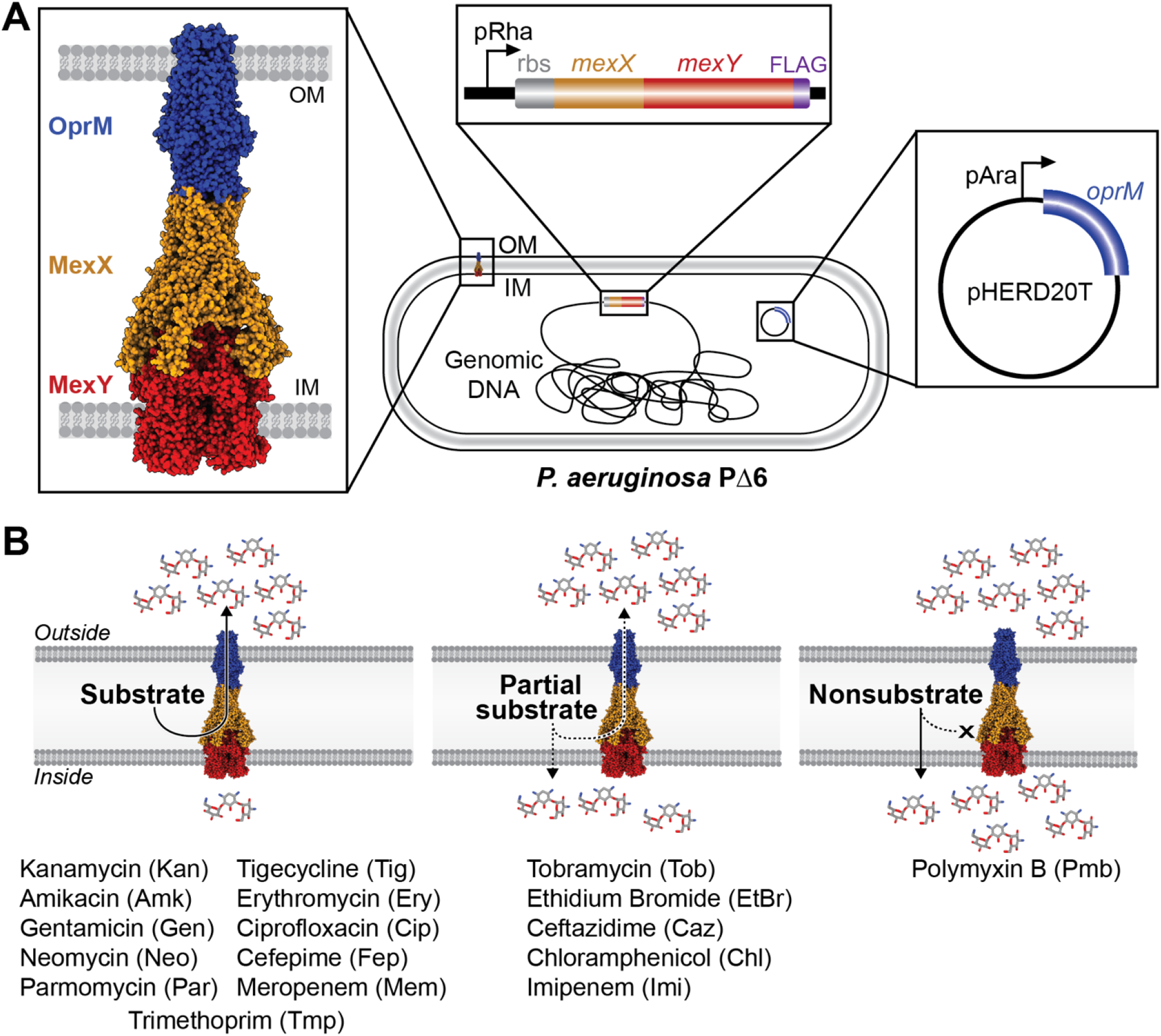
Overview of the MexXY-OprM induction system and substrate classes. *A*, Homology model *(left*) of the tripartite complex containing outer membrane (OM) protein OprM (blue), periplasmic adaptor protein MexX (orange), and inner membrane (IM) transporter MexY (red). Cartoon schematic of the transposon system containing the rbs and PAO1 *mexXY* with FLAG-tag (purple) under the control of a rhamnose promoter in the PΔ6 genomic DNA (*center*) and *oprM* under an arabinose promoter located on the pHERD20T plasmid (*right*). ***B***, Schematic representation of the three substrate classes (substrate, partial substrate, and nonsubstrate) with MexXY-OprM located in the cell envelope. Antibiotics identified in each class are given below.

To overcome these limitations, we aimed to identify substrate preference for PAO1 MexXY through its controlled induction from a genomic chromosomal insertion in a multi-deletion *P. aeruginosa* background, PΔ6 (PAO1 Δ*mexAB-oprM, mexCD-oprJ, mexEF-oprN, mexJKL, mexXY, triABC*)^17, 18^. PΔ6 was selected for these studies as the six deleted pumps are the major contributors to intrinsic antibiotic resistance in *P. aeruginosa* and have overlapping substrate profiles which would otherwise complicate MexXY-OprM substrate identification. Additionally, in the absence of these additional efflux systems, measurements of minimum inhibitory concentrations (MIC) can reliably serve as a proxy for MexXY-OprM efflux activity for each potential substrate. The native *mexXY* operon and ribosome binding site (rbs) were PCR amplified from *P. aeruginosa* PAO1 genomic DNA and cloned into the miniTn7 vector, pJM220^19^, under a rhamnose inducible promoter (resulting in pTn7XY), and a FLAG tag-encoding sequence subsequently appended 3’ of *mexY* (resulting in pTn7XY-FLAG; **Tables S1** and **S2**). Using transposon insertion, pTn7XY-FLAG and helper plasmid, pTNS3, were electroporated into PΔ6. As outlined previously^20^, insertion of rbs-*mexXY* and removal of the gentamicin resistance marker using pFLP were confirmed using colony PCR (resulting in strain LK17). To complement the absent OMP in LK17 (due to deletion of *mexAB-oprM* in PΔ6), *oprM* was amplified from PAO1 genomic DNA and cloned into the arabinose-inducible complementation plasmid, pHERD20T^21^, and electroporated into both PΔ6 and LK17 resulting in strains LK36 and LK21, respectively (**Fig. 1A**, and **Tables S1** and **S2**). This approach allows for MexXY overexpression in the same isogenic genetic background and without the need for non-native chimeric operon construction with the OMP, which may result in polar effects on transcription.

Optimal inducer concentrations for MexXY (rhamnose) and OprM (arabinose) were empirically determined, with 1% wt/vol of each inducer determined to show the largest increase in resistance without affecting cell growth (**Fig. S1** and **Table S3**). To quantify efflux-mediated resistance for a panel of chemically and functionally diverse antimicrobials, we measured the MIC of each potential substrate as previously described^22, 23^, in the absence or presence of inducer (i.e., rhamnose and/or arabinose) in all strains (**Fig. 2** and **Table S4**). Western blot analysis was performed on whole cell lysate after seven hours of rhamnose induction to confirm MexY expression in LK21 (**Fig. S2**).

**Fig. 2.**
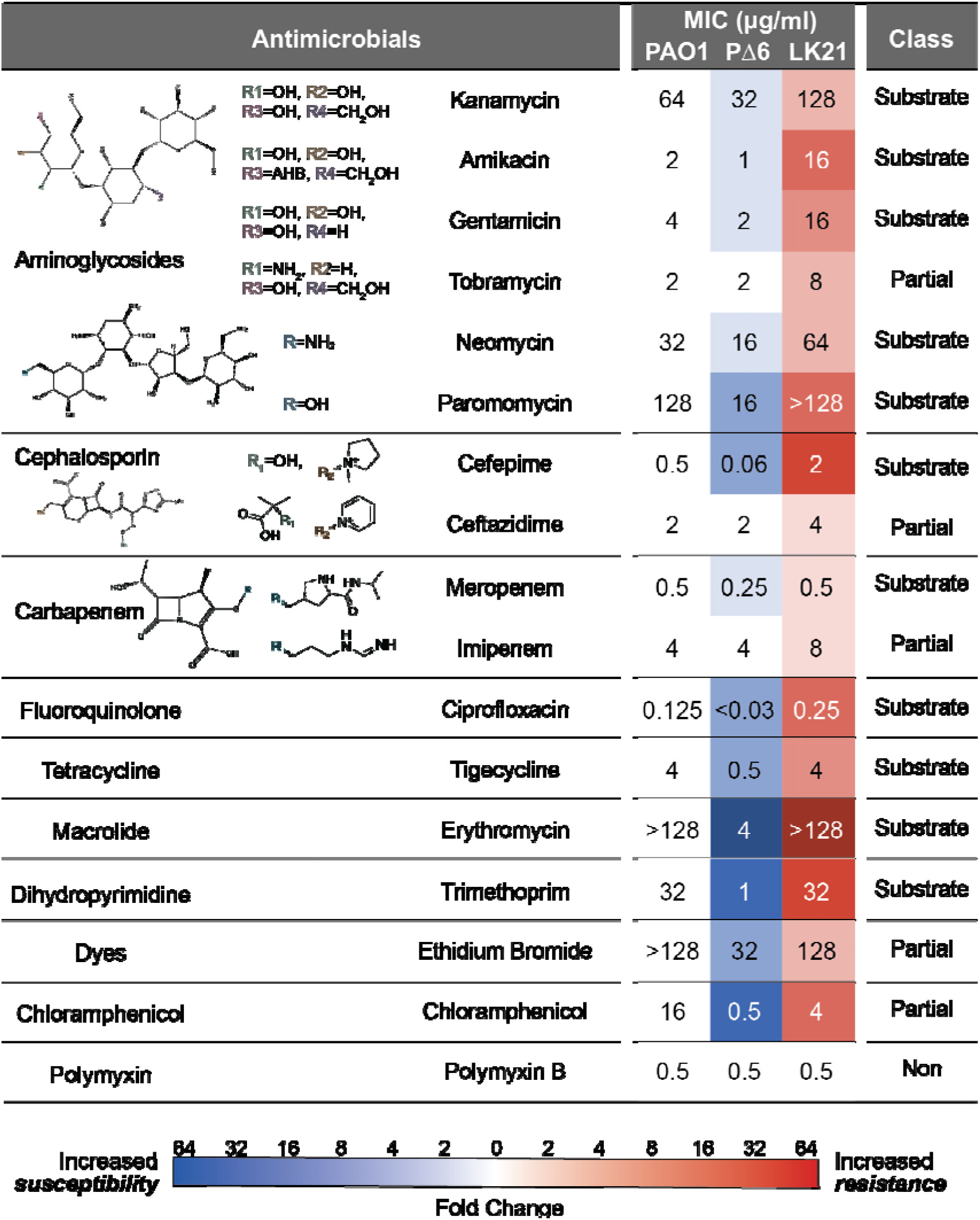
Substrate profile of the MexXY-OprM efflux pump in the *P. aeruginosa* PΔ6 expression system. *Left*, antibiotic families and the individual antibiotics tested; where two or more antibiotics within the same family were tested, differences in their chemical structure are indicated. *Center*, average MICs (µg/mL) for 10 classes of antibiotics in *P. aeruginosa* PAO1, PΔ6, and LK21 (PΔ6::mexXY^PAO1^,pHERD-oprM^PAO1^) in the presence of 1% wt/vol arabinose and rhamnose are shown numerically overlayed on a heatmap indicating fold-change compared to parent strain (PΔ6 is compared to PAO1 and LK21 is compared to PΔ6). Blue is increase in susceptibility (reduced efflux) and red is increase in resistance (increased efflux). *Right*, the resulting substrate classification for each antibiotic is indicated as substrate (Substrate), partial substrate (Partial) or nonsubstrate (Non).

Deletion of the six efflux systems in PΔ6 resulted in increased sensitivity to all antibiotics tested with the exception of tobramycin (Tob), ceftazidime (Caz), imipenem (Imi), and polymyxin B (Pmb), which are weak or non-substrates for RND-type efflux pumps in *Pseudomonas*^24, 25^. Substrates with the highest change in susceptibility (up to 64-fold) were Ery, Chl, and trimethoprim (Tmp), confirming that the deleted RND efflux systems are collectively a major contributor to resistance to these antimicrobials in *P. aeruginosa* PAO1 (**Fig. 2**). Interestingly, the use of arabinose and rhamnose sugars antagonized action of aminoglycosides in both PAO1 and PΔ6, as well as Ery, cefepime (Fep), and Chl in PAO1 (**Table S4**). We speculate that because aminoglycosides are modified sugars, the presence of additional sugars in the media may regulate other aminoglycoside entry/efflux transporters^26^.

Overexpression of MexXY-OprM in PΔ6 allowed for robust efflux of aminoglycosides, ciprofloxacin (Cip), Fep, tigecycline (Tig), Ery, meropenem (Mem), and Tmp, for which MICs were restored to or exceeded wild-type PAO1 levels. We refer to these antimicrobials as substrates of MexXY-OprM. These analyses also revealed Tmp to be a previously undefined substrate for MexXY-OprM (**Fig. 1B, Fig. 2** and **Table S4**). We further classified the tested antibiotics into two additional categories: ‘partial substrates’ or ‘nonsubstrates’. We define partial substrates as either effluxed antimicrobials for which resistance levels did not change in PΔ6 compared to PAO1 but for which resistance increased moderately in the presence of MexXY-OprM expression, or when resistance levels were decreased in PΔ6 but overexpression of MexXY alone could not restore the resistance observed in PAO1 (**Fig. 1B**). For example, resistance to EtBr and Chl increased but was not restored to the level observed for PAO1 upon MexXY-OprM overexpression, suggesting that additional RND systems or mechanisms are partly or primarily responsible for resistance to these substrates. Additionally, the MICs for Tob, Caz, and Imi did not change upon deletion of the six efflux systems in PΔ6 but were increased upon overexpression of MexXY-OprM (**Table S4**). These data suggest that these antimicrobials are not preferred MexXY-OprM substrates but may be exported when the pump is overexpressed, or that the overexpression of MexXY-OprM itself leads to physiological changes that impact resistance to these antimicrobials. Furthermore, as these compounds are members of antibiotic families with substrates, it is possible that the small differences in chemical substituents may result in loss of efflux recognition (**Fig. 2**). The only antibiotic tested whose resistance was unaffected by the deletion or complementation of MexXY-OprM was Pmb which is thus classified as a nonsubstrate.

Overall, these studies identified Tmp as a MexXY-OprM substrate and the potential for MexXY-OprM to contribute to Caz and Imi resistance when overexpressed, a phenomenon typically observed in CF clinical isolates^27, 28^. As our system allows for the overexpression of MexXY-OprM, without significantly impacting growth, the increased threshold allows identification of subtle differences within substrate classes (e.g., aminoglycoside, cephalosporins, and carbapenems). Furthermore, this system can enable future studies on transporter-adaptor pairs with exchangeable plasmid-encoded OMPs allowing for detailed analyses of OMP impact on substrate selectivity. As efflux-mediated resistance is a major contributor to multidrug resistance, and therefore therapeutic failures, such approaches to better understand efflux substrate specificity as well as the impact of overexpression on ‘nonsubstrate’ antibiotics are of vital importance.

## Acknowledgements

This work was supported by the National Institutes of Health (NIH) National Institute for Allergy and Infectious Disease award R01 AI185192 (to G.L.C.) and NIH National Institute for General Medical Sciences NRSA Pre-doctoral Fellowship F31 GM143891 (to L.G.K.).

