## Supplemental Materials for "Determination of *Pseudomonas aeruginosa* MexXY-OprM substrate profile in a major efflux knockout system reveals distinct antibiotic substrate classes"


**Running Title:** MexXY-OprM substrate profile


**Key Words:** Efflux, *Pseudomonas,* antimicrobial susceptibility, antimicrobial resistance, resistance-nodulation-division (RND)


This file contains:

**Supplemental Figures:**


**Figure S1**. Rhamnose induction assay for MexXY expression and cell viability
**Figure S2**. MexY expression in strain LK21


**Supplemental Tables:**


**Table S1**. Strains and plasmids used in this study
**Table S2**. DNA primers used in this study
**Table S3**. Gentamicin minimum inhibitory concentration (MIC) in LK21 in the presence of variable rhamnose and arabinose inducer concentrations for MexXY and OprM expression
**Table S4**. Antimicrobial susceptibility data for *P. aeruginosa* PAO1 and PΔ6 expressing MexXY, OprM, or the whole efflux pump MexXY-OprM.

**Supplemental Figures**

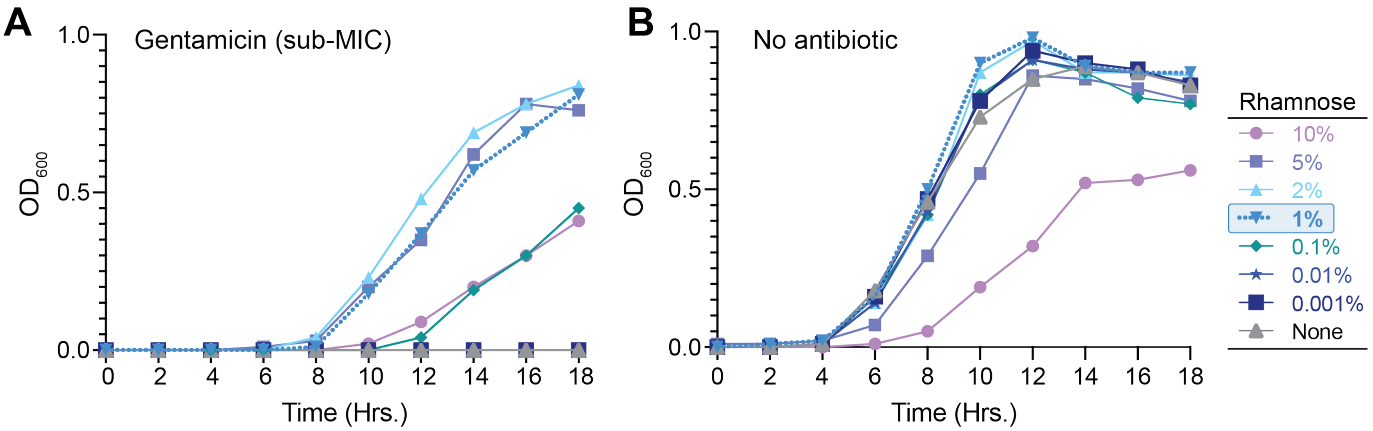


**Fig. S1. Rhamnose induction assay for MexXY expression and cell viability.** Optical density at 600 nm ($OD_{600}$) of strain LK21 grown over 18 hours in cation-adjusted Mueller-Hinton broth supplemented with 1% arabinose ***A***, at the greatest subinhibitory concentration of gentamicin (8 µg/mL) and ***B***, in the absence of antibiotic at variable concentrations (0.001-10%) of rhamnose. The no rhamnose control is shown in grey triangle and the selected inducer concentration (1%) is highlighted with the blue dotted line.

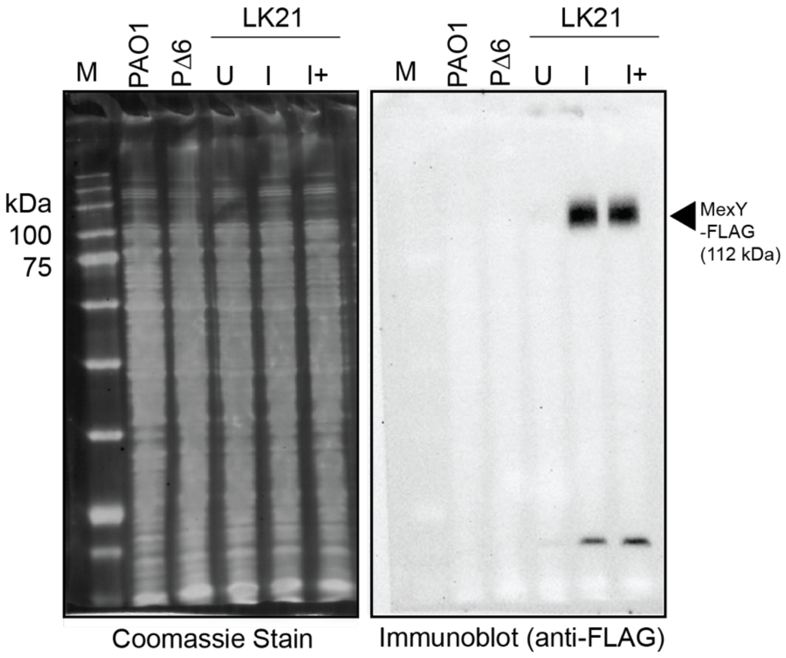


**Fig. S2. MexY expression in strain LK21.** Coomassie stained SDS-PAGE gel (*left*) and HRP-conjugated anti-FLAG (ABclonal) immunoblot used to detect the C-terminal MexY FLAG (*right*). Lanes shown are whole cell lysate for PAO1, PΔ6, and LK21 with no inducer (U), MexXY induced with 1% wt/vol rhamnose (I), or MexXY and OprM induced with 1% wt/vol rhamnose and arabinose (I+). MexY-FLAG (112 kDa) migrated the appropriate size shown by protein marker (M).

### Supplemental Tables

#### Table S1. Strains and plasmids used in this study.

| Strain | Genotype | Reference |
|---|---|---|
| PAO1 | *P. aeruginosa* laboratory strain | Goldberg Lab |
| PΔ6 | PAO1 Δ*mexAB-oprM*, Δ*mexCD-oprJ*, Δ*mexEF-oprN*, Δ*mexJKL*, Δ*mexXY*, Δ*triABC* | 1,2 |
| LK17 | PΔ6::$mexXY^{PAO1}$ | This study |
| LK21 | PΔ6::$P_{rha}$-rbs-$mexXY^{PAO1}$, pHERD20T-$P_{BAD}$-$oprM^{PAO1}$ | This study |
| LK36 | PΔ6, pHERD20T- $P_{BAD}$-$oprM^{PAO1}$ | This study |
| **Plasmid** | **Genotype** | **Reference** |
| pJM220 | pUC18-miniTn7T-gm-rhaSR-$P_{rha}$BAD | 3 |
| pTn7XY | pUC18-miniTn7T-gm-rhaSR-PrhaBAD-*rbs_*$mexXY^{PAO1}$ | This study |
| pTn7XY-FLAG | pUC18-miniTn7T-gm-rhaSR-PrhaBAD-*rbs_*$mexXY^{PAO1}$-*FLAG* | This study |
| pHERD20T | pUCP20T-araC-$P_{BAD}$ | 4 |
| pHD20TM | pUCP20T-araC-$P_{BAD}$-$oprM^{PAO1}$ | This study |

#### Table S2. DNA primers used in this study.

| Primer name | Sequence (5'-3') |
|---|---|
| PAO1_HindIII_RBS*mexX-F* | GCGAAGCTTTGAACGTCCTCACAAGGGAAAG |
| PAO1_ApaI_*mexY-R* | GCGGGGCCCTCAGGCTTGCTCCGTG |
| PAO1_EcoRI_*oprM-F* | GCGGAATTCATGAAACGGTCCTTCCTTTC |
| PAO1_HindIII_*oprM-R* | GCGAAGCTTTCAAGCCTGGGGATCTTC |
| MexY_FLAG-F | GACTACAAGGACGACGATGACAAGTGACTCGCGAAGGCC |
| MexY_FLAG-R | GGCTTGCTCCGTG |

#### Table S3. Gentamicin minimum inhibitory concentration (MIC) in LK21 in the presence of variable rhamnose and arabinose inducer concentrations for MexXY and OprM expression.

| | | Gentamicin MIC (µg/mL) | | | | | | | |
|---|---|---|---|---|---|---|---|---|---|
| MexXY induction[a] | ***% Rha*** | ***10*** | ***5*** | ***2*** | ***1*** | ***0.1*** | ***0.01*** | ***0.001*** | ***0*** |
| | | 16 | 16 | 16 | **16** | 16 | 8 | 8 | 2 |
| OprM induction[b] | ***%Ara*** | ***4*** | ***2*** | ***1*** | ***0.5*** | ***0.25*** | ***0.125*** | ***0.06*** | ***0*** |
| | | 16 | 16 | **16** | 8 | 8 | 8 | 4 | 4 |

[a]Arabinose (OprM inducer) maintained at fixed concentration of 1%.

[a]Rhamnose (MexXY inducer) maintained at fixed concentration of 1%.

**Table S4. Antimicrobial susceptibility data for *P. aeruginosa* PAO1 and P∆6 expressing MexXY, OprM, or the whole efflux pump MexXY-OprM.**

| | | Minimum inhibitory concentrations (µg/mL)[a] | | | | | | |
|---|---|---|---|---|---|---|---|---|
| | | **PAO1** | | **P∆6** | | **LK17** | **LK36** | **LK21** |
| | Genotype | WT | | ∆ABM/ CDJ/ EFN/ JKL/ XY/ triABC | | P∆6 + XY | P∆6 + M | P∆6 + XYM |
| *Inducer* | *rha* | - | + | - | + | + | - | + |
| | *ara* | - | + | - | + | - | + | + |
| Aminoglycosides | Kan | 64 | 64 | 16 | 32 | 32 | 32 | 128 |
| | Amk | 1 | 2 | 0.5 | 1 | 2 | 1 | 16 |
| | Gen | 2 | 4 | 0.5 | 2 | 2 | 1 | 16 |
| | Tob | 1 | 2 | 1 | 2 | 2 | 2 | 8 |
| | Neo | 16 | 32 | 8 | 16 | 16 | 16 | 64 |
| | Par | 64 | 128 | 8 | 16 | 16 | 32 | >128 |
| Tetracycline | Tgc | 4 | 4 | 0.5 | 0.5 | 1 | 0.5 | 4 |
| Macrolide | Ery | 128 | >128 | 4 | 4 | 16 | 16 | >128 |
| Fluoroquinolone | Cip | 0.25 | 0.125 | <0.03 | <0.03 | <0.03 | <0.03 | 0.25 |
| Dye | EtBr | >128 | >128 | 32 | 32 | 32 | 32 | 128 |
| Cephalosporin | Fep | 0.25 | 0.5 | 0.06 | 0.06 | 0.06 | 0.125 | 2 |
| | Caz | 2 | 2 | 2 | 2 | 2 | 2 | 4 |
| Chloramphenicol | Chl | 8 | 16 | 0.5 | 0.5 | 1 | 1 | 4 |
| Carbapenem | Mem | 0.5 | 0.5 | 0.25 | 0.25 | 0.06 | 0.25 | 0.5 |
| | Imi | 4 | 4 | 4 | 4 | 4 | 4 | 8 |
| Trimethoprim | Tmp | 32 | 32 | 0.5 | 1 | 0.5 | 1 | 32 |
| Polymyxins | Pmb | 0.5 | 0.5 | 0.5 | 0.5 | 0.5 | 0.5 | 0.5 |

[a]Antimicrobials: kanamycin (Kan); amikacin (Amk); gentamicin (Gen); tobramycin (Tob); neomycin (Neo); paromomycin (Par); tigecycline (Tig); erythromycin (Ery); ciprofloxacin (Cip); ethidium bromide (EtBr); cefepime (Fep); ceftazidime (Caz); chloramphenicol (Chl); (Mem); imipenem (Imi); polymyxin B (Pmb); trimethoprim (Tmp)
